# Effects of Δ*motY* mutations on motility behavior of *Pseudomonas aeruginosa* chimeric periplasmic stator variants

**DOI:** 10.1101/2025.11.06.686970

**Authors:** Nicola Ferguson, Mahika Desai, Joscelyn Flores Silva, Hoa Nguyen, Bruce Rodenborn, Orrin Shindell, Frank Healy

**Affiliations:** Department of Biology, Trinity University, San Antonio, Texas, 78212 USA; Department of Mathematics, Trinity University, San Antonio, Texas, 78212 USA; Physics Program, Centre College of Kentucky, Danville, Kentucky, 40422 USA; Department of Physics and Astronomy, Trinity University, San Antonio, Texas, 78212 USA

**Keywords:** Flagellar motility, stators, MotY, swimming, swarming, periplasmic binding domain

## Abstract

*Pseudomonas aeruginosa* utilizes dual flagellar stator systems for motility; and dual stator bacteria possess auxiliary flagellar rotor ring components in their periplasm. In *P. aeruginosa* MotAB and MotCD comprise the dual stator system and MotY is the auxiliary periplasmic ring component. We investigated motility of strains and their isogenic Δ*motY* derivatives which expressed chimeric MotB/MotD periplasmic domains and characterized differences in motility behaviors. We found in general motility is severely impaired in strains carrying *motY* deletions and which express C-terminal periplasmic regions of MotD, compared with strains expressing MotB C-terminal counterparts. Motility in soft agar is slightly increased in strains expressing N-terminal MotB transmembrane domains and MotD C-terminal periplasmic plug, PGB, and extensions, but motility is severely impaired in Δ*motY* strains. Addition of the extended 24-residue C-terminus of MotB to the C-terminus of MotD does not significantly affect either motility or compensate for the deleterious effect of Δ*motY* mutation, but does significantly increase motility in *motY+* backgrounds. The soft agar motility results for organisms with wild type MotAB stators was not different from those with MotAB stators carrying 24 residue MotB C-terminal deletions; however, motility of these mutants was significantly lower in Δ*motY* mutants compared to Δ*motY* mutants expressing wild type MotAB stators. We discuss contributions of stator functional domains to motility and the effects of Δ*motY* deletion. Lastly, we speculate on possible mechanistic roles for the two stator plug types based on thermodynamic considerations related to differences in composition of the hydrophobic surfaces of the two amphipathic helices.

**IMPORTANCE:** *Pseudomonas aeruginosa* uses two torque-generating stators, MotAB and MotCD, to drive flagellar rotation. Dual stator bacteria have additional rotor components; in *P. aeruginosa*, this component has been identified as MotY. This study investigates the interaction between the C-terminal plug and periplasmic regions of the MotB and MotD components of the MotAB and MotCD stator complexes with MotY. Motility assays of periplasmic chimeric strains expressing variants with MotB and MotD C-terminal plug and peptidoglycan binding domains reveal an enhanced sensitivity of MotD C-terminus to MotY deletions. These data suggest that critical interactions, either direct or indirect, must take place between the basal body MotY ring complex and MotD periplasmic regions for proper function of the MotCD stator in flagellar motility.

## INTRODUCTION

Many bacteria use flagella to propel themselves through fluid environments and across surfaces (1). The flagellum is a macromolecular reversible rotary nanomachine consisting of an extended filament propeller connected to a basal body motor through a flexible universal joint hook. Extensive genetic, biochemical and structural studies of the Salmonella basal body motor assembly reveal a central axial rod passing through a series of rings embedded in the cell surface. These include the C (cytoplasmic), MS (membrane/supramembrane), P (peptidoglycan), and L (lipopolysaccharide) rings. (2).

Flagella are powered by ion flow, with filament rotational energy originating from ion transduction through membrane-diffusible stators. The stator consists of two proteins present in a heptameric A_5_B_2_ stoichiometry, referred to as MotAB, MotCD, PomAB, etc. (3, 4). Motor rotation occurs through the association of stator complexes with the C ring. The *Escherichia coli* flagellar motors recruit up to a maximum of about a dozen stators, in a load-dependent manner, to drive motor rotation (5, 6). MotA contains four transmembrane (TM) helices, and each of five MotA peptides associates into a truncated pentagonal cone, which contacts the C ring. The A_5_ pentamer surrounds the N-terminal TM helices of the central MotB dimer and extends into the periplasm. The MotB TM helix is followed by a plug helix, a disordered linker region and C-terminal peptidoglycan binding domain (PGB). A critical aspartate residue located within the MotB TM region is required for ion transport (corresponding to D33 in both *S. enterica* and *P. aeruginosa*) (7). The plug helix prevents ion flow through freely diffusing stators not associated with the rotor (8). Comparison of plugged and unplugged states in the *Campylobacter jejuni* MotAB complex reveal a variety of conformational changes that take place to create a transmembrane ion channel and initiate proton flow. Concomitant with motor recruitment of a stator unit, MotB PGB self-dimerizes to anchor the membrane-diffusible unit in association with the motor (9–11).

Many bacteria, such as *Salmonella* sp., utilize a single torque-generating stator complex, some species use dual stator systems and do so in different ways and contexts (12, 13). *Vibrio* and *Shewanella* spp. have Na+ dependent PomAB stators in addition to H+ dependent MotAB stator (14, 15). *Pseudomonas aeruginosa* expresses two H+ driven stators, MotAB and MotCD (C_5_D_2_) (16, 17). Different motility phenotypes are observed between *P aeruginosa* strains expressing one stator or the other. *P. aeruginosa* PA01 Δ*motCD* strains swim faster than Δ*motAB* strains, while MotCD stators are required for motility in viscous environments and along surfaces (16). In *P. aeruginosa* PA14, swimming speeds are similar for both stators in low viscosity fluid media; however, in high viscosity media, Δ*motCD* mutants are non-motile, indicating that MotCD is required for motility in high viscosity environments (17). MotAB and MotCD stator dynamics studies show MotCD-driven rotational speeds are lower than those of MotAB (18).

While MotB and MotD exhibit overall sequence and structural similarities, there are notable differences in the periplasmic domains of these proteins. The linker region between the helix plug and PGB is 17 residues longer in MotB than in MotD; and the C-terminal extension beyond the PGB in MotB is 38 residues longer than that of MotD (Figure 1).

**Figure 1.**
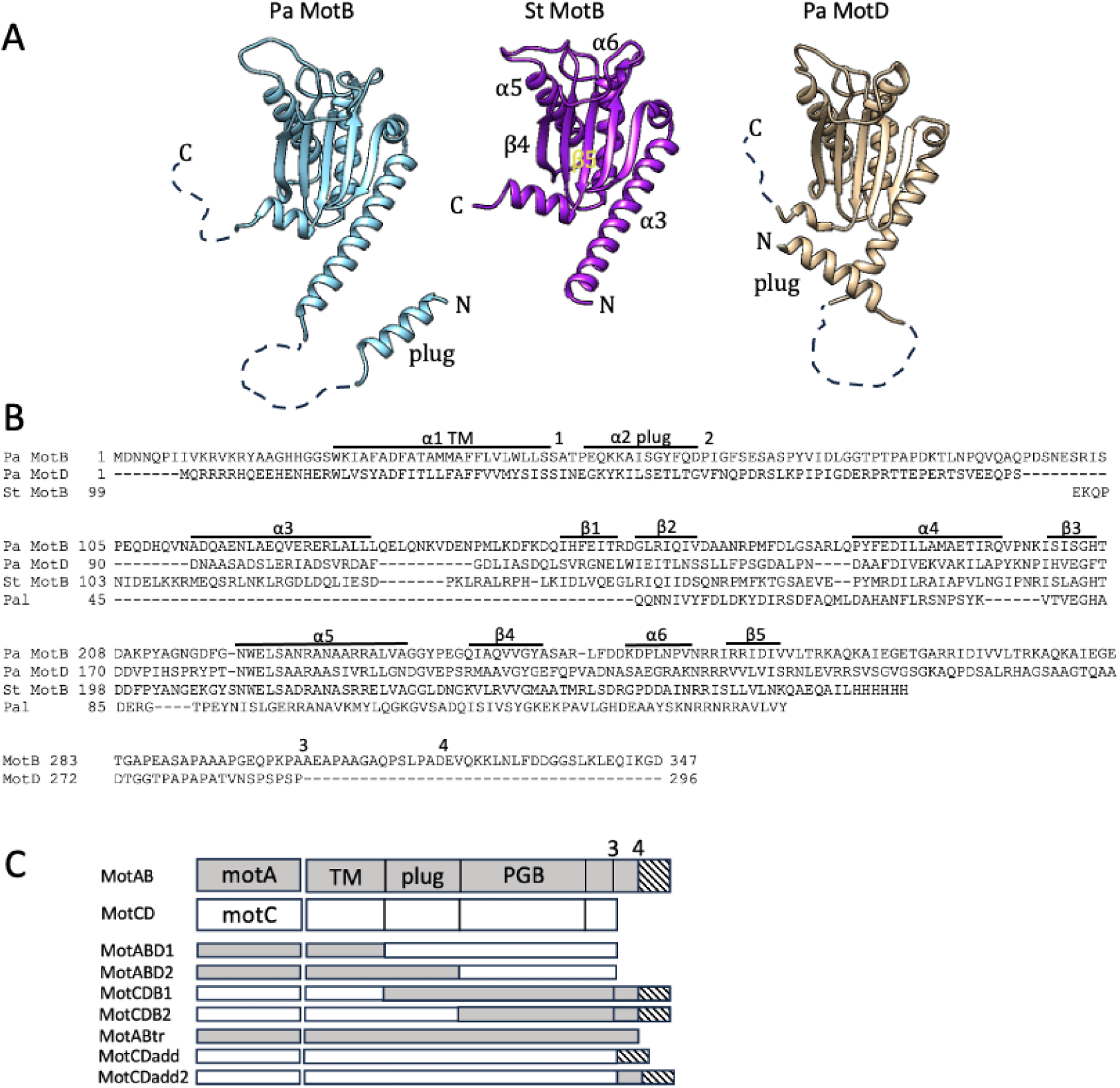
Structural and sequence features of *P. aeruginosa* MotB/MotD periplasmic domains and chimeric strains. (A) Comparison of S. typhimurium MotB (St MotB) periplasmic structure (PDB 2ZVY (35)) with *P. aeruginosa* MotB and MotD periplasmic structure models. N-terminal transmembrane domains are omitted; periplasmic linker sequence between plug helix and helix 3, and C-terminal extensions beyond PGB are shown as dashed lines. Select secondary structural feature labels of St MotB are shown. (B) Structure-based alignment of P. aeruginosa MotB and MotD C-terminal periplasmic region using *Salmonella typhimurium* MotB PDB 2Z0V. (C) Composition of inducible Tn7-integrated chimeric stator constructs. Components originating from MotAB are shown in gray, components originating from MotCD are shown in white.

In addition to the canonical rotor C, MS, L and P rings, electron microscopy and electron cryo-tomography (ECT) studies reveal accessory features in the periplasmic regions of basal body complexes of bacteria with dual stator systems, such as *P. aeruginosa*. A structure situated below the LP ring system was identified in *V. alginolyticus* and named the T ring, comprising MotX and MotY (19). Analysis of *V. alginolyticus* Δ*motX* and Δ*motY* mutants shows that the PomAB stator system requires MotX and MotY for incorporation or stabilization of the PomAB stator in the motor. Structural studies have demonstrated that MotY is required for MotX recruitment to the basal body, and MotY prevents breakdown of MotX (20, 21). ECT studies and bioinformatic analysis of dual stator system bacteria show a correlation between the presence of dual stator systems, elaborations around the basal body P ring region, and the occurrence of *motY* homologs in dual stator organism genomes. ECT-derived basal body structures from *Shewanella oneidensis*, *Legionella pneumophila*, and *P. aeruginosa* show elaborated *motY*-encoded structures below the LP ring system, variously described as T rings or decorated P rings. Previous genetic studies in *P. aeruginosa* PA01 demonstrated that MotY was required for MotCD-dependent motility (16). In bead-based assays using flagellar filament stubs to study MotY:stator interaction dynamics, FRET analysis of MotY donor-MotB/MotD acceptor energy transfer shows strong molecular interactions between MotY and both MotB and MotD (18, 22). Significant differences in motor rotation speeds and switching rates between wild type *P. aeruginosa* and isogenic Δ*motY* strains are observed; and a 30% decrease in pole-localized stator fluorescence is seen relative to wild type strain. Swimming plate assays showed that Δ*motAB* Δ*motY* strains were essentially non-motile, similar to Δ*motB* Δ*motCD* strains.

Collectively, these results show MotAB and MotCD participate in critical interactions with T ring MotY. MotY plays a role in regulating motor torque and switching rates, though it remains unclear how the *P. aeruginosa* MotAB/CD dual stator system might participate in this process. And while MotY also plays a role in some aspect of stator recruitment and/or stabilization, the manner through which MotY:stator interactions take place is unknown.

To investigate stator:MotY interactions in this dual stator system, we constructed stator variant strains expressing inducible chimeric alleles of *motB* and *motD*, where periplasmic domains of one subunit are exchanged with its counterpart. These strains and their isogenic Δ*motY* derivatives were then used in soft agar, liquid medium, and swarming assays to investigate interactions of MotY with periplasmic regions of MotB and MotD.

## RESULTS

To investigate periplasmic interactions between MotY and MotAB and MotCD complexes and their effect on cell motility, a Δ*motAB*Δ*motCD* stator mutant was used as host for Tn7-mediated integration of rhamnose-inducible chimeric stator constructs. The stator variant strains were designed to compare the impact on motility phenotype of C-terminal regions MotB and MotD in *motY+/-* genetic backgrounds. Strain construction terminology is illustrated in Figure 1C, showing strains MotABD1, MotABD2, MotCDB1, and Mot CDB2, with swapped chimeric regions; strain MotABtr, with truncated MotB C-terminal extension; and strains MotCDadd and MotCDadd2, which express motD fused to a 24-(MotCDadd) or 38-(MotCDadd2) residue C-terminal tail of MotB. Growth of all strains was similar to wild type PA01 in L broth and VBMM citrate media.

### Differential soft agar motility responses

The results of soft agar motility experiments are presented first in Figures 2 and 3. Based on diameters of soft agar motility zones, strains could be divided into six groups: 1) weakly motile; 2) MotAB-type; 3) MotCD-type; and 4) dual-type. A fifth group consisted of ABD1 and CDB1Δ*motY*, with motility zones that were consistently intermediate between MotAB-type and MotCD-type. (compare Figure 2 panels F and G; and panels F and K). A sixth category consisted of strain MotCDadd and MotCDadd2, which displayed soft agar swimming behaviors that were intermediate between MotCD-type and wild-type. Motility of wild type PA01 and strains expressing single stators, and their Δ*motY* counterparts is shown in Figure 2 panels A-D; motility of PA14 Δ*motAB* and Δ*motCD* strains is shown in Figure 2 panel E. Motilities of the Tn7::*motAB* and Tn7::*motCD* single stator constructs were similar to PA01 strains expressing single stators (compare Figure 2 panels B and J, and Figure 3 columns 3 and 11; and Figure 2 panels C and F, and Figure 3 columns 5 and 10). The Δ*motAB*Δ*motCD* host strain carrying the control Tn7 vector was non-motile (compare Figure 2 panels D, F, and J).

**Figure 2.**
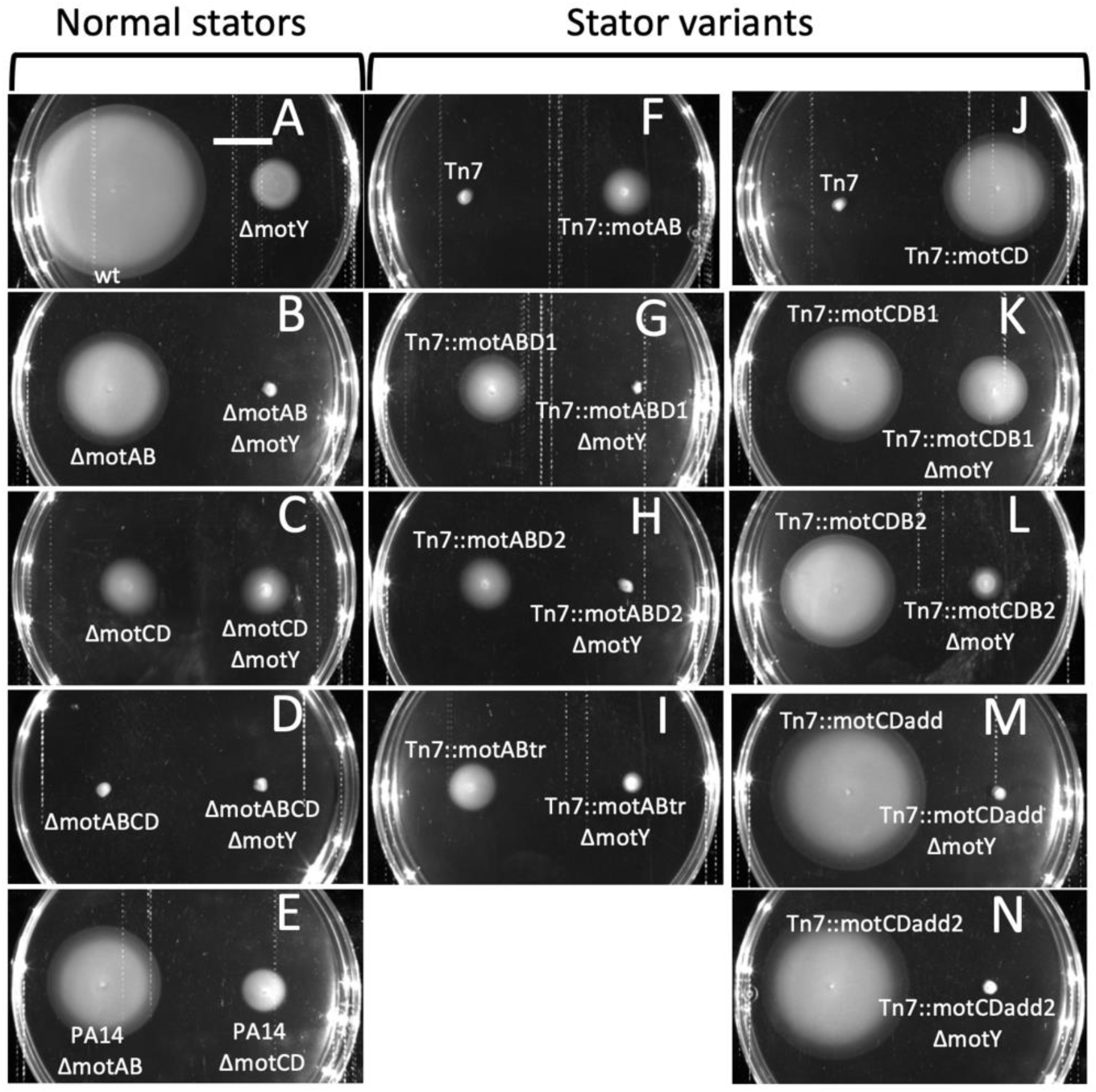
Motility behavior of MotB/MotD chimeric stator strains and ΔmotY derivatives on 0.3% motility agar. Pinpoint inocula were incubated for 22 hr at 37C. Panels are arranged vertically: left column, normal stator configuration (zero, one, or two stators); center column, chimeric MotAB strains withTn7:motAB control shown in panel F; right column, chimeric MotCD strains with Tn7:motCD control shown in panel J. Panel A bar = 15 mm.

**Figure 3.**
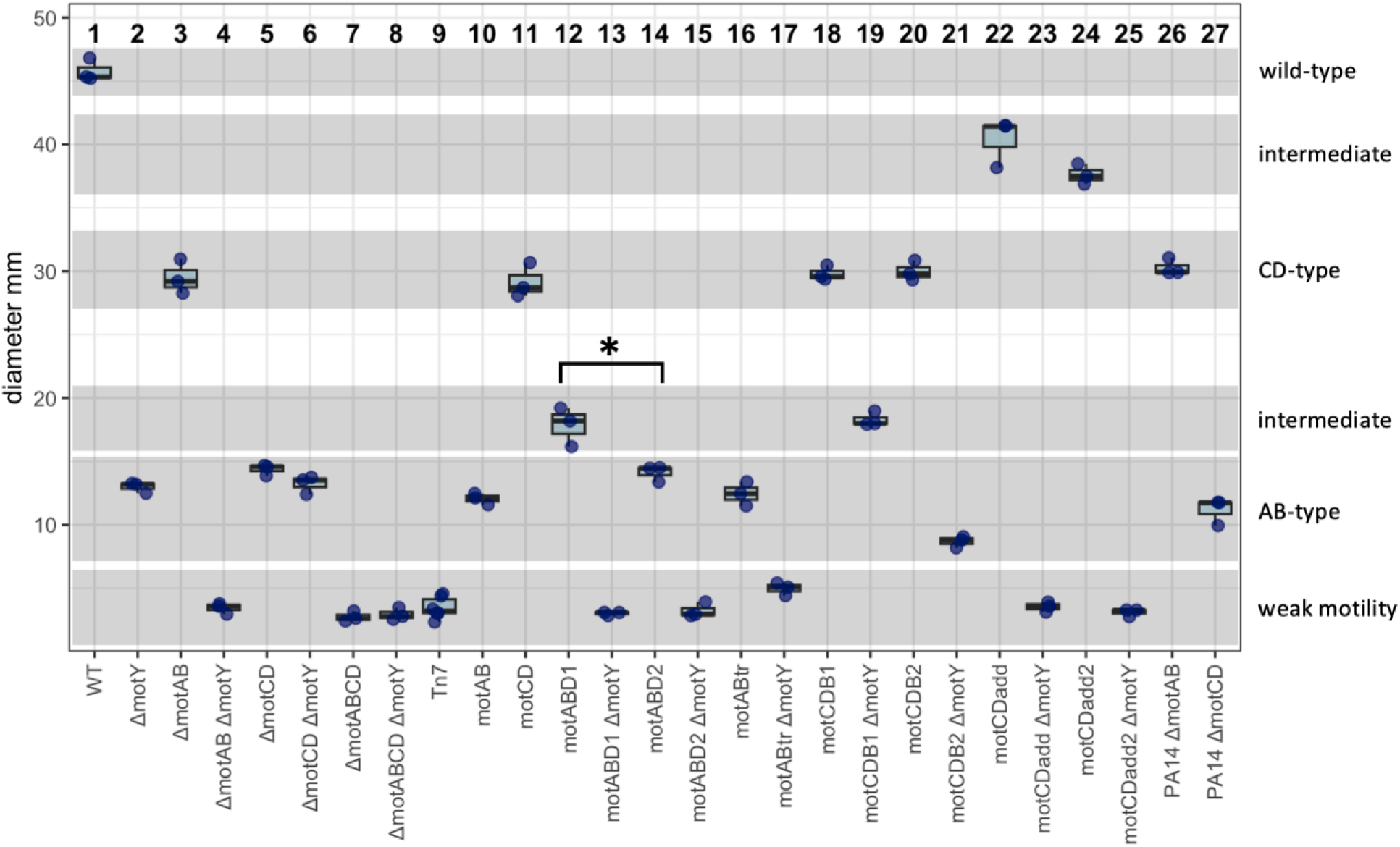
Soft agar motility analysis of stator variant strains. Experiment was done as described in Figure 2. ImageJ tools were used to measure diameters of radial colony expansion. Values represent means ± standard deviation (n=3). Significant differences were assessed using the Student’s t-test assuming equal variances; *t* = 3.86, *p* = 0.018.

The effect on motility of the *motY* deletion between the two stators was striking, and similar to previously reported findings (16). The Δ*motY* effect on motility was severe in all strains expressing MotD periplasmic regions compared with strains expressing MotB periplasmic regions (Figure 2 compare Δ*motY* mutants in panels B, G, and H with Δ*motY* mutants in panels C, K, and L). Motility of Δ*motY* mutants expressing the single MotCD stator complex is severely impaired, compared to the motility effect in cells expressing the single MotAB stator in Δ*motY* backgrounds (compare Figure 2 panels B and C; Figure 3 compare columns 3 and 4 versus columns 5 and 6).

We next compared motility behavior of the stator variants and their Δ*motY* derivatives. While the strain MotABD1 motility was greater than strain MotAB, the deleterious effect of Δ*motY* on strain MotABD1, which expressed the MotD periplasmic domain (including MotD plug), was considerably greater than the effect of Δ*motY* on motility of the MotAB strain (compare Figure 2 panels C and G; Figure 3 compare columns 5 and 6 versus columns 12 and 13). Also, the motility of strain MotABD2, expressing the MotD periplasmic linker and C-terminus, was similar to that of the MotAB strain, and motility in this variant was more adversely affected in Δ*motY* mutants compared to MotAB strains (compare Figure 2 panels C and H; Figure 3 compare columns 5 and 6 versus columns 14 and 15). It is notable from these observations that the severity of the Δ*motY* motility impairment is greater in chimeric strains expressing MotD plug domains.

The effect of the Δ*motY* mutation on motility was less severe in stator variant strains carrying MotB periplasmic regions. The motility of strain MotCDB1, expressing MotB plug and C-terminus, was similar to that of the MotCD strain, but the deleterious effect of Δ*motY* in the MotCD strain was considerably greater than the effect of the Δ*motY* mutation in the MotCDB1 strain (Figure 2 compare panels B and K; Figure 3 compare columns 3 and 4 versus columns 18 and 19). We also see increased sensitivity of Δ*motY* mutants to motility impairment in chimeric strains expressing the MotD plug helix domain in MotCDB1 and MotCBD2 strains. The Δ*motY* motility impairment of MotCDB1 (MotB plug) is less severe than the Δ*motY* impairment effect on MotCDB2 (MotD plug).

The effect of Δ*motY* mutation on MotAB stator variant with truncated C-terminus is shown in Figure 2 panel I. In the MotABtr strain, with a deletion of 24 residues from the MotB C-terminus, motility values were similar to those of MotAB and MotAB Δ*motY* strains. However, the motility of the MotABtr Δ*motY* strain was considerably impaired relative to MotAB, MotABtr and MotAB Δ*motY* strains (compare Figure 2 panels C and I; Figure 3 compare columns 5 and 6 versus columns 16 and 17).

The motility of MotCDadd, which MotD carries the 24 C-terminal residues of MotB, was consistently greater than the motility of the MotCD strain; however, the effect of Δ*motY* mutation was similar for MotCD and MotCDadd (compare Figure 2 panels B and M, and Figure 3 columns 3 and 22). The motility of the MotCDadd2 strain carrying the extended 38-residue MotB C-terminus was slightly but reproducibly lower than MotCDadd. Isogenic Δ*motY* derivatives of both MotCDadd and MotCDadd2 strains were non-motile in the soft agar swimming assay.

### MotB/MotD periplasmic domain and motY effects on swimming in liquid media and swarming motility

Swimming trajectories for each of the strains were recorded in liquid media and the data are presented in Figure 4. Trajectories were distributed over a broad range for MotAB, Δ*motCD* Δ*motY*, and MotABD1 (compare Figure 4 columns 4 – 6). Swimming was significantly impaired in the MotABD2 strain (MotB plug and MotD C-terminus). Compared with their MotB counterparts, swimming speeds were low for all strains expressing MotD N-termini, but differences in the effect of Δ*motY* were observed. MotCD Δ*motY,* MotABD1 Δ*motY*, and MotABD2 Δ*motY* strains were non-motile, indicating that MotD C-terminal domain requires MotY for motility in liquid media. Swimming speeds for MotCDB1 and MotCDB2 strains were similar to their isogenic Δ*motY* counterparts. Swimming speeds of MotABtr and MotABtrΔ*motY* strains were similar to the wild type, indicating that the MotB C-terminal extension is not required for swimming in liquid media. The MotCDadd Δ*motY* strain (with partial 24-residue MotB C-terminal extension) was non-motile. Motility was not completely abolished, but was severely impaired in MotCDadd2 Δ*motY* strains (with full 38-residue MotB C-terminal extension).

**Figure 4.**
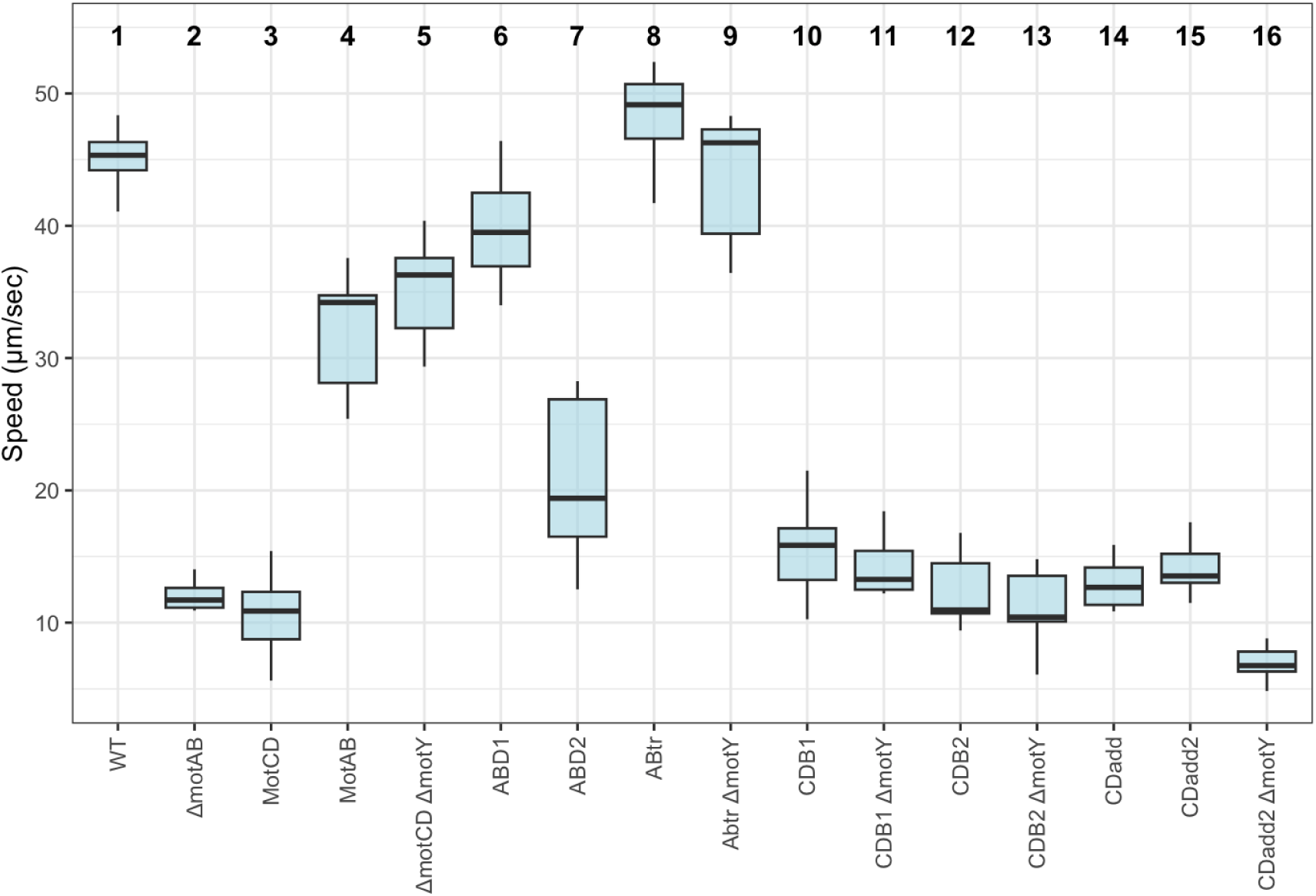
Summary swimming data in motility medium. Trajectories of exponential phase cultures of cells grown on motility medium were acquired on a Zeiss Axioscop II using darkfield optics and analyzed using ImageJ plug-ins. At least ten trajectories were acquired and analyzed for each strain. Stator variant strains ΔmotAB ΔmotY, Tn7::motCDadd ΔmotY, Tn7::ABD1 ΔmotY, Tn7::ABD2 ΔmotY were non-motile under these conditions.

Swarming behavior of stator variants is presented in Figure 5. Wild-type strain PA01 displayed the simple circular swarming pattern typical for the strain (strain PA14 swarming is shown for comparison, panels M-O) (23). Swarming is severely impaired or abolished in Δ*motY* strains. MotAB variants with MotB N-termini were non-motile. Swarming behavior in MotCDB1 strains expressing MotD N-terminus with MotB plug and extended C-terminus was similar to strains expressing MotCD; however, swarming was impaired in MotCDB2 strains expressing MotD N-terminus and plug region with MotB C-terminus. Swarming motility in MotCDadd strains was slightly better than MotCD strains.

**Figure 5.**
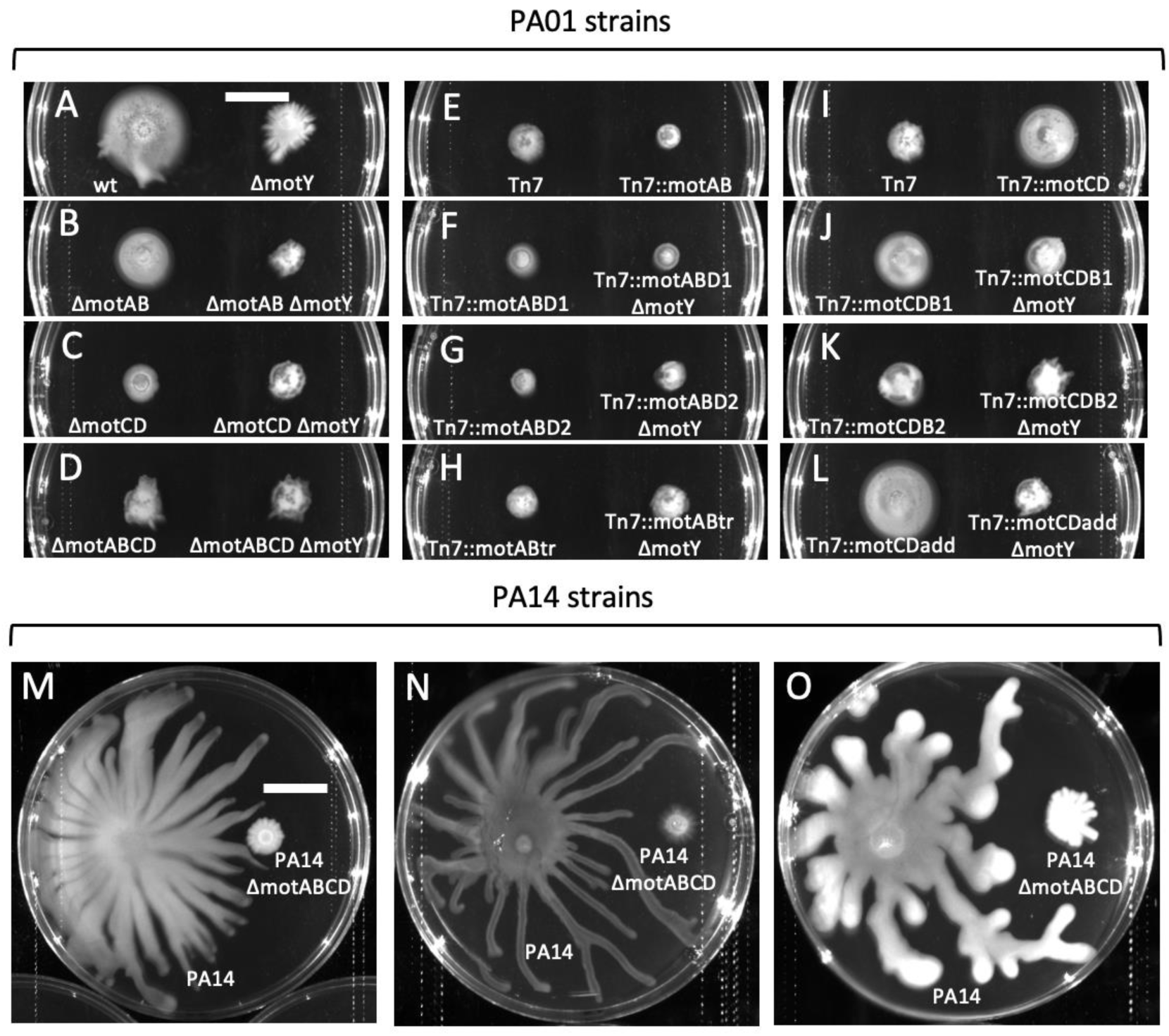
Swarming motility of stator variants. Top panels, *P. aeruginosa* PA01 strains, arranged vertically: left column, normal stators (zero, one or two stators) and Δ*motY* derivatives; center column, MotAB and MotAB Δ*motY* chimeric stator strains; left column, MotCD and MotCD Δ*motY* chimeric strains. These panels show growth on M9 glutamate swarming media (45); M9 aspartate and casamino acids swarm media (23) produced similar results. Plates were incubated for about 22 hr at 37C and imaged. Bottom three panels M – O, swarm behavior of *P. aeruginosa* PA14 wild-type and dual stator Δ*motABCD* strains on M, M9 glutamate; N, M9 aspartate; and O, casamino acids swarm media.

## DISCUSSION

In *P. aeruginosa* and some other bacteria, the torque forces that drive flagellar motility are provided by two distinct stator complexes. Additionally, these dual stator organisms possess auxiliary flagellar ring components, such as MotY, in the periplasmic space. To explore periplasmic flagellar motor-stator interactions in the *P. aeruginosa* dual stator system, we created strains expressing rhamnose-inducible chimeric variants of MotB and MotD periplasmic domains and their isogenic Δ*motY* derivatives. Numerous studies have been done using organisms expressing stators of various chimeric composition (e.g., 16, 24–27). In this work we have selected the model dual stator organism to investigate the roles of MotB/MotD periplasmic functional domains in motility and their interactions, either direct or indirect, with MotY. This approach may provide new insights into mechanistic aspects of stator function regarding the specific roles played by MotB/MotD periplasmic domains in flagellar dynamics. Indeed, it is likely that modular domains in dual stator organisms would undergo an optimization suited to and directed towards optimal activity under the conditions in which it operates (e.g., surfaces, fluid viscosity, Na+ or H+, etc.). Thus, characterizing the contributions of functional domains to motility behavior could inform our understanding of flagellar dynamics.

Motility in soft agar assays was greater, and swimming speeds were generally higher, in strains expressing MotB with MotD plug and MotD C-terminus (MotABD1). This effect was reversed in soft agar assays with strains that expressed MotB N-termini including MotB plug, with MotD C-termini (MotABD2). In the liquid swimming assay, swimming speeds for strains expressing MotB N-termini including the MotB plug, with MotD C-termini (MotABD2) were lower than that of normal MotAB strains. On the other hand, substitution of MotB plugs in chimeric strains expressing MotD N-termini made no observable differences in motility. Motility was similar for strains expressing MotCD stators with either MotD or MotB plugs.

The behavior of strains MotABD2 and MotCDB2, which expressed chimeric N-terminal PGB, was similar to strains expressing non-chimeric PGB. On the other hand, in both cases the motility behavior of isogenic Δ*motY* derivatives was significantly impaired. However, no significant differences in motility were observed between strains expressing MotAB and their isogenic Δ*motY* derivatives. This observation could be explained by proposing that the extended length of the MotB C-terminus compared to MotD contributes to stator function in a MotY-independent manner. MotB is 51 residues longer than MotD, most of this extended length (38 residues) occurs at the C-terminus. Our findings support the idea that this disordered end contributes to some aspect of MotAB flagellar activity and does not require MotY. An extended 38-residue C-terminus would have a theoretical maximum length of about 13 nm (28); this could provide opportunities for critical interactions across a periplasmic space with a width on the order of 10 – 20 nm (29). The data suggest MotAB activity has an absolute requirement for either MotY or the extended C-terminus, as motility of MotABtr Δ*motY* strains was severely impaired. In our analysis of MotB/MotD sequences, the MotB homolog in dual stator organisms is typically longer than its MotD counterpart.

We uncovered a plug effect in both soft agar and liquid swimming motility tests. Motility of MotABD1, a MotAB stator with MotD plug, was better than MotABD2, a MotAB stator with MotB plug. Motility of the ABD1 strain was also consistently higher than strains expressing normal MotAB complexes. Interestingly, the motilities of strains MotCDB1 and MotCDB2 were both similar to one another and to strains expressing normal MotCD stators, but the effect of the Δ*motY* mutation on motility of MotCDB2, expressing the MotD plug, was more severe than the effect of the Δ*motY* mutation on MotCDB1, which expresses the MotB plug. Thus, somehow the impact of the Δ*motY* deletion is greater in strains expressing MotD plugs.

Genetic and biochemical studies of the *Escherichia coli* MotB stator component show that the plug adopts an amphipathic alpha helical structure (8). In the plugged state, the MotB dimer can function in trans, with the ion channel for MotB_1_ closed by the MotB_2_ plug. The helix is proposed to embed itself in the hydrophobic membrane surface along a hydrophobic face of the amphipathic helix, thus obstructing ion flow through a MotA/MotB transmembrane channel. Electron cryo-tomography studies have identified a likely channel which, in the plugged state, is obstructed by a conserved MotA phenylalanine residue, F186 in the *C. jejuni* structure (4); a corresponding site in the *P. aeruginosa* model is F210. Structures of unplugged complexes show the phenylalanine side chain position rotated in such a way as to leave the transmembrane channel open. In its activated state, it has been proposed that the MotB plugs are displaced from the membrane and form an association between their respective hydrophobic faces (8). We analyzed the properties of plug helices from both stator forms of several dual stator organisms. Helical wheel plot analysis was used to identify and characterize differences between the MotB and MotD plugs, and results are shown in Figure 6. The amphipathic hydrophobic helical surfaces and opposing hydrophilic faces of MotB and MotD are depicted alongside the *S. typhimurium* MotB structure (panel A). By comparison, the helices are largely unremarkable except for the occurrence of a basic lysine residue on the MotB hydrophobic surface (panel B).

**Figure 6.**
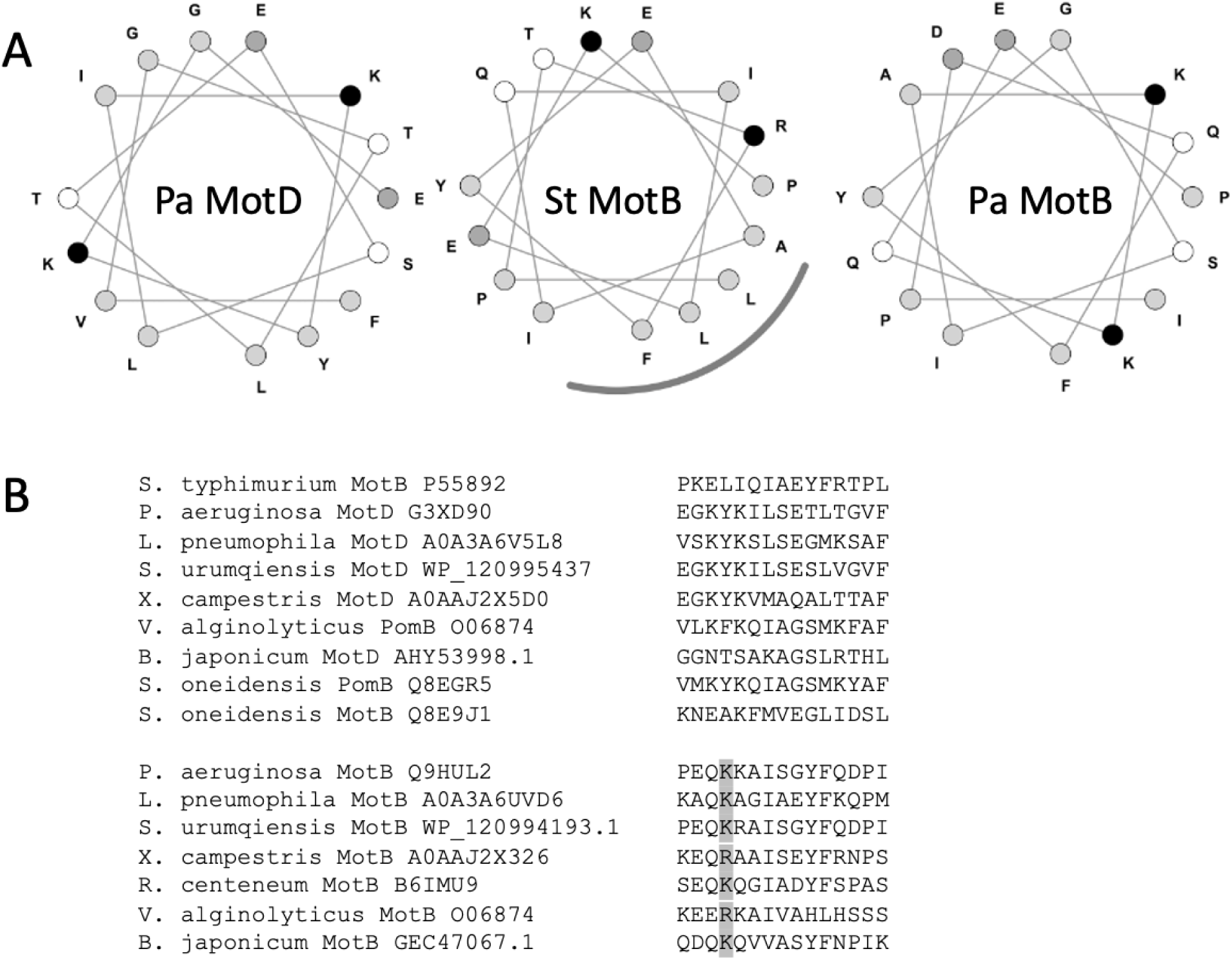
Helical wheel plot analysis and sequence alignments of amphipathic plug helices for dual stator system organisms. (A) Helical wheel plots for *P. aeruginosa* MotD (Pa MotD), *S. typhimurium* MotB (St MotB), and *P. aeruginosa* MotB (Pa MotB). Polar uncharged residues are white, nonpolar residues are light gray, acidic residue are dark gray, and basic residues are black. Hydrophobic helical face is indicated with gray arc along lower right quadrant of St MotB wheel. (B) Sequence alignments of MotB/MotD stators for dual stator organisms. *S. typhimurium*-type stators have nonpolar residues in hydrophobic surface position four, whereas *P. aeruginosa* MotB-type stators have Arg or Lys residues in position four (highlighted).

A recent comprehensive structural and phylogenetic analysis of bacterial stators identified two structural evolutionary lineages, flagellar ion transporters (FIT) and generic ion transporters (GIT) (30). Characterization of 107 FITs revealed several shared features among flagellar stator proteins and absent in GITs: 1) extended cytoplasmic domain with at least four helices comprising the torque-generating interface; 2) a square four-helix fold transmembrane domain in the MotA subunit; 3) MotB subunit plug and linker domains; and 4) expanded peptidoglycan binding domain. FITs diverged structurally into TGI4 and TGI5 groups, based upon the presence of four (TGI4) or five (TGI5) helices within the stator torque-generating interface. TGI5 stators, e.g., *E. coli* MotAB, are mostly restricted to members of the Pseudomonadota. TGI4 stators include Na+-driven systems such as *V. alginolyticus* PomAB. Our limited analysis of dual stator plug composition revealed that the MotB-type stators carry lysine residues in the fourth position of the plug helix (Figure 6, panel A). Logo analysis of the 107 stator dataset show high bit scores for lysine or arginine at this position in MotB-type stators (30). This strong correlation across multiply divergent stator evolutionary histories and gene transfer phenomena would suggest that the occurrence of a basic residue on the hydrophobic surface plays some specific role in flagellar activity; and conversely, the specific functionality of plug helices with hydrophobic surfaces characterized by strictly nonpolar residues may be dependent on the surface being exclusively hydrophobic.

Since we find stator plugs with nonpolar and basic residues at position four in the stator complexes of dual stator organisms, we propose that in dual stator organisms, the composition of the plug at position four might play an important role in some mechanistic aspect of flagellar activity. We explored this proposal by addressing bioenergetic properties of the two helix types. We used the neural network-based model PMIpred (31) to estimate free energies of interaction of the nonpolar and lysine-containing amphipathic helices. In addition to providing estimates for hydrophobicity and hydrophobic moment, the model also assesses propensities for interactions with curved surfaces. Helices with lower free energies which bind curved surfaces are “sensors”, whereas helices with stronger energies (“binders”) exhibit non-selectivity. PMIpred analysis of plug helices is shown in Figure 7. Plugs with lysine occupying position four typically have lower free energies with a curvature-sensing component; plugs with nonpolar residues at position four do not exhibit curvature selectivity.

**Figure 7.**
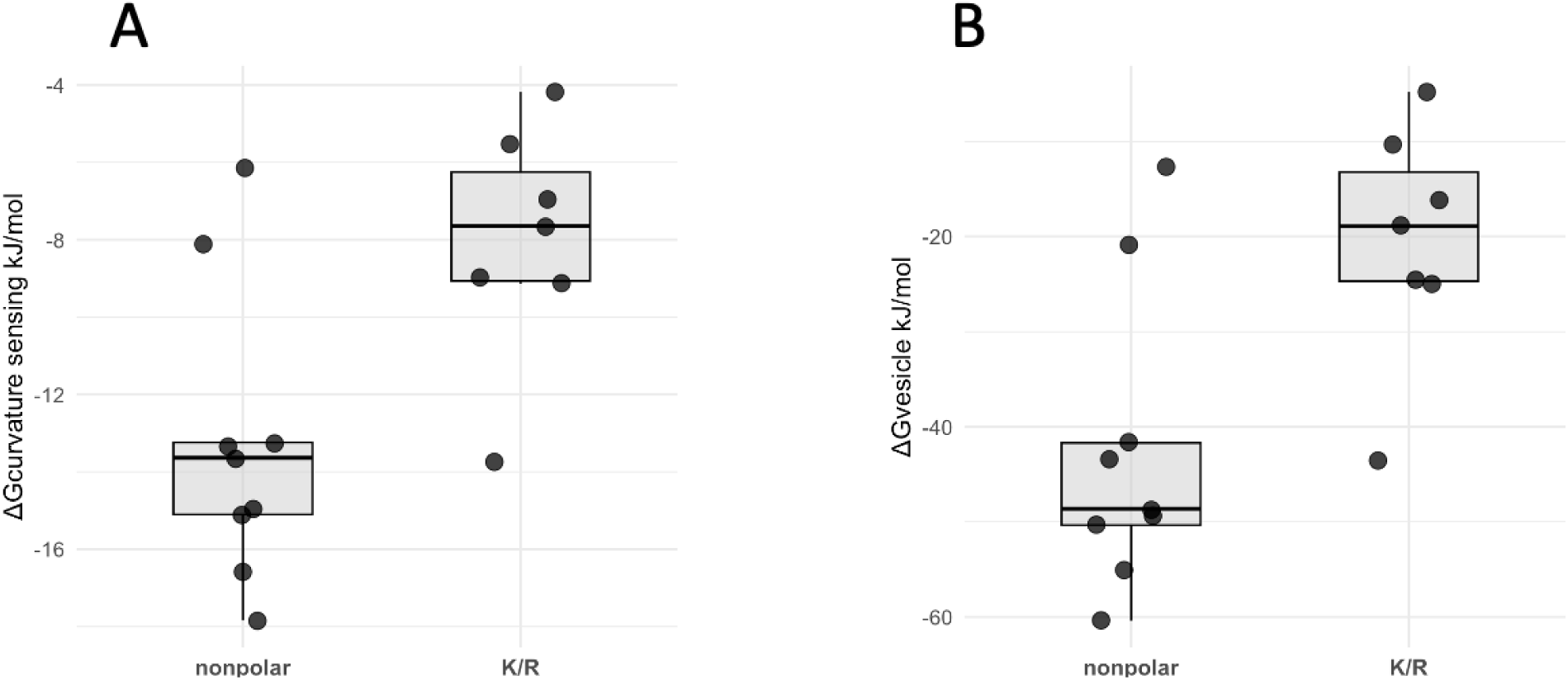
PMIpred modeling analysis of MotB/D plug helices. The protein-membrane interaction neural network model PMIpred (31) was used to predict free energies of membrane-associative interactions for MotB/D proteins listed in Figure 6 based on amino acid type at helix position four: either nonpolar at position four; or a lysine or arginine at position four, K/R). (A) Predicted curvature-sensing free energy, ΔΔF_adj, adjusted for charge effects. (B) Predicted membrane-spanning free energy, ΔF_sm(R=50), which estimates the free energy for a vesicle with radius = 50 nm.

We are intrigued by the idea that differences in the thermodynamic properties of the two plug types, and the observation of “plug effects” in our motility behavior assays, would suggest that some mechanistic aspect of the differential behavior of stators in dual stator bacteria is governed by the interfacial membrane dynamics of the two stator types. For example, at the channel activation stage, could plug type be sensitive to differences in local membrane properties? Or, following activation, could stator turnover dynamics be regulated by changes in membrane properties? The divergence of plug types based on structural changes which significantly affect thermodynamic properties could represent an interesting example of an evolutionary mechanistic specialization of periplasmic amphipathic helices.

## MATERIALS AND METHODS

### Bacterial strains, plasmids and media

Strains and plasmids used in this work are given in Table 1. *Pseudomonas aeruginosa* strain PA01 (JJHO MPA01 re-sequenced) was a gift from Joe Harrison; PA14 strains were a gift from George O’Toole; *Escherichia coli* strain S17.1 and plasmid pTNS2 (Addgene plasmid #64968) were gifts from Herbert Schweizer. Bacterial strains were routinely cultured on lysogeny broth (per liter: peptone, 10 g; yeast extract, 5 g; sodium chloride, 5 g). Liquid motility media contained (per liter) 5 g peptone, and 5 g sodium chloride; soft agar motility plates consisted of motility media plus 0.3% agar. Swarm media was prepared with casein hydrolysate (23) and M8 glucose medium with 0.05% aspartate or glutamate (45), amended with 0.55% agar. VBMM was prepared as described by Choi and Schweizer (46); NSLB was prepared with 300 mM sucrose as described by Hmelo et. al. (47). Antibiotics were included where appropriate at the following concentrations: ampicillin, 100 ug/mL (*E. coli*); gentamicin, 10 ug/mL (*E. coli*) and 30 ug/mL (*P. aeruginosa*). Stator expression in pJM220 Tn7 insertion strains was induced with addition of rhamnose to 0.3 mM in plate media, and 3 mM in liquid media.

**Table 1.** Bacterial strains and plasmids used in this study.

| Strain/plasmid | Genotype/properties | Source or reference |
| --- | --- | --- |
| PA01 |  |  |
| L30 | JJH0: wild type Illumina re-sequenced MPA01 | Joe Harrison (49) |
| L32 | JJH0 ΔmotY | This study |
| L36 | JJH0 ΔmotAB | This study |
| L37 | JJH0 ΔmotAB ΔmotY | This study |
| L42 | JJH0 ΔmotCD | This study |
| L43 | JJH0 ΔmotCD ΔmotY | This study |
| L48 | JJH0 ΔmotABCD | This study |
| L53 | JJH0 ΔmotABCD ΔmotY | This study |
| L73 | JJH0 ΔmotABCD Tn7 | This study |
| L74 | JJH0 ΔmotABCD Tn7::motAB | This study |
| L75 | JJH0 ΔmotABCD Tn7::motCD | This study |
| L54 | JJH0 ΔmotABCD Tn7::motABD1 | This study |
| L62 | JJH0 ΔmotABCD Tn7::motABD1 ΔmotY | This study |
| L56 | JJH0 ΔmotABCD Tn7::motABD2 | This study |
| L64 | JJH0 ΔmotABCD Tn7::motABD2 ΔmotY | This study |
| L58 | JJH0 ΔmotABCD Tn7::motCDB1 | This study |
| L66 | JJH0 ΔmotABCD Tn7::mot CDB1 ΔmotY | This study |
| L60 | JJH0 ΔmotABCD Tn7::mot CDB2 | This study |
| L68 | JJH0 ΔmotABCD Tn7::mot CDB2 ΔmotY | This study |
| M1 | JJH0 ΔmotABCD Tn7::motABtr (motBΔ323-347) | This study |
| M13 | JJH0 ΔmotABCD Tn7::motABtr (motBΔ323-347) ΔmotY | This study |
| M7 | JJH0 ΔmotABCD Tn7::motCDadd (motD + motB323-347) | This study |
| M19 | JJH0 ΔmotABCD Tn7::motCDadd (motD + motB323-347) ΔmotY | This study |
| M64 | JJH0 ΔmotABCD Tn7::motCDadd2 (motD + motB310-347) | This study |
| M66 | JJH0 ΔmotABCD Tn7::motCDadd (motD + motB310-347) ΔmotY | This study |
| PA14 |  |  |
| I9 | Wild type | George O'Toole |
| I10 | ΔmotAB | George O'Toole |
| I11 | ΔmotCD | George O'Toole |
| I12 | ΔmotABCD | George O'Toole |
| *E. coli* S17.1 | thi pro hsdR hsdM recA RP4 tra λpir | Simon et. al. (50) |
| *E. coli* NEBα | $\Delta$(lacZ)M15 gyrA96 recA1 relA1 endA1 hsdR17 | NEB |
| Plasmids |  |  |
| pEX18Gm | ColE1 Gm<sup>r</sup> oriT sacB | Hoang et. al. (51) |
| pJM220 | pUC18T-miniTn7T- Gm<sup>r</sup>-rhaSR-PrhaBAD | addgene #110559 (48) |
| pTNS2 | bla R6Kγ ori tnsABCD | Choi et. al. (52) |

### Construction of strains

P aeruginosa PA01 *ΔmotAB* and *ΔmotCD* single stator and *ΔmotABΔmotCD* deletion mutants were constructed using previously published methods (47). One kb PCR amplification products flanking the *motAB* and *motCD* loci were joined in plasmid pEX18Gm, and plasmids were used to transform *E. coli* S17.1 (supplementary table).Transformants and cultures of *P. aeruginosa* PA01 were collected on filters for conjugal mating as described (47). Merodiploid PA01 transconjugants were selected on VBMM medium containing gentamicin; and single-crossover deletion mutants were selected on NSLB medium containing sucrose. Silicon-based platform synthesis was used to generate stator *motAB*, *motCD* and stator variant constructs (Twist Bioscience). Figure 1 illustrates the stator variant constructs. Ribosome binding site sequence 5’CCGTTTACAGAGAGGAGAACG was added to the 5’ ends of *motA* and *motC* open reading frames. Variant constructs carrying *motA* retained the *motAB* intergenic region. ABD1 carries the N-terminal TM helix domain through MotB Ser47 and the remaining C-terminal sequence is that of MotD, from the plug region Ile 41 through the remainder of the MotD sequence. ABD2 contains the MotB TM and MotB plug through Gln 61, and the remainder of the sequence is that of MotD from Gln 59 to the end of the sequence. Variant constructs carrying *motC* retained the *motCD* intergenic region. CDB1 carries the N-terminal MotD TM through Ser 40, and the remaining sequence is that of MotB, including plug (Pro 50 – Gln 61), linker, PGB and extended C-terminus. CDB2 contains the MotD TM and MotD plug, and the remainder of the expressed protein is that of MotB, including linker, PGB, and extended C-terminus. CDadd expresses MotCD with the additional 24 C-terminal residues of MotB (from Asp 324 to Asp 347); CDadd2 expresses the 38-residue MotB C-terminal extension. ABtr expresses MotAB with the MotB C-terminal 24 residues removed. All stator variant constructs were cloned into the PstI site of pJM220 under transcriptional control of the PrhaBAD promoter (48). Strain L48 (*ΔmotABΔmotCD*) was used as host for Tn7-mediated transposition of pJM220 derivatives carrying stator constructs using electroporation with helper plasmid pTNS2 (46).

### Motility assays

For soft agar motility assays, toothpick inoculum from a fresh overnight plate culture or 3 uL of an overnight L broth culture were transferred to plates and plates were incubated for about 20 hr at 37C before imaging. Swarm plates were inoculated as previously described (23) and imaged after about 24 hr of growth at 37C. Images were processed using ImageJ. Overnight or exponential phase cultures in liquid motility medium were used for swimming assays. Wet mounts were prepared using a small volume of cell culture and observed using dark field optics with a Zeiss Axioscop 2 equipped with a camera (Thorlabs 1501M-USB). Swimming trajectories were recorded and analyzed using ImageJ particle tracking plug-in.

## ACKNOWLEDGMENTS

We thank Joe Harrison and George O’Toole for providing strains. This research was supported by National Science Foundation grants PHY-2210609 and PHY-2210610.

---

**Supplementary Table.**
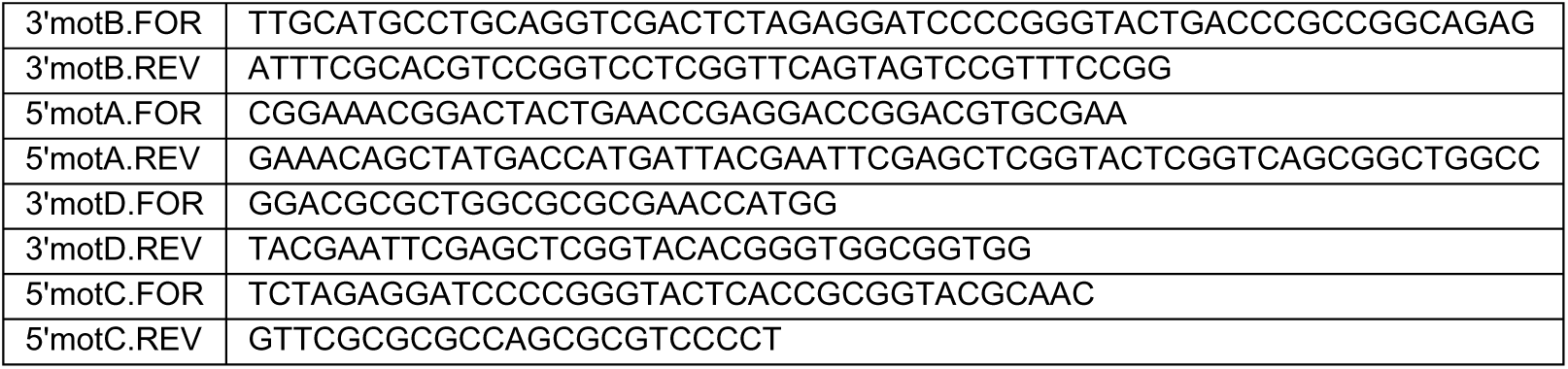
Oligonucleotides used for construction of PA01 stator mutants.

**Supplementary Table 1.**
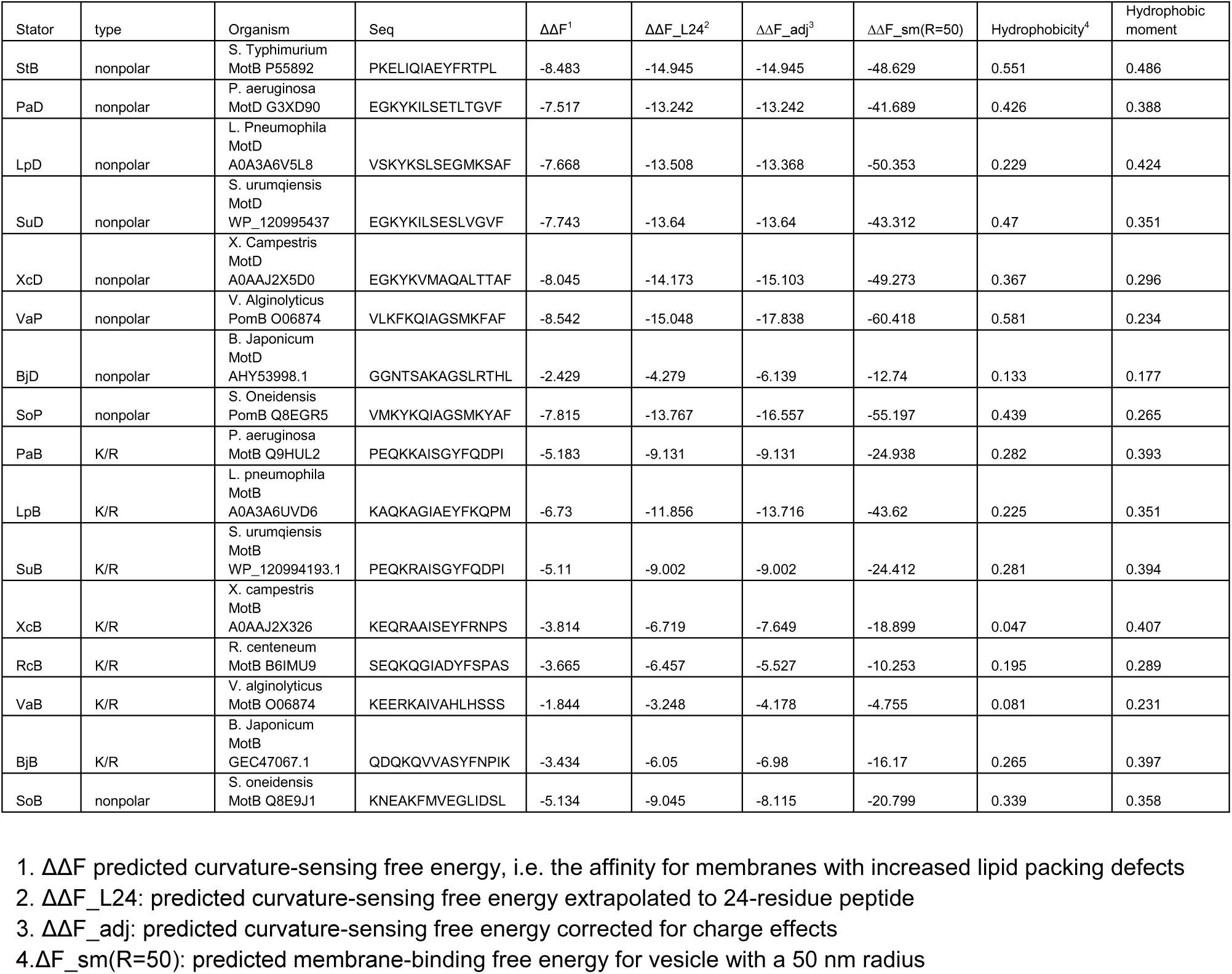
PMIpred free energy values for amphipathic stator plug helices.

